# Retrospective inference as a form of bounded rationality, and its beneficial influence on learning

**DOI:** 10.1101/569574

**Authors:** Thomas HB FitzGerald, Will D. Penny, Heidi M Bonnici, Rick A Adams

## Abstract

Probabilistic models of cognition typically assume that agents make inferences about current states by combining new sensory information with fixed beliefs about the past, an approach known as Bayesian filtering. This is computationally parsimonious, but, in general, leads to suboptimal beliefs about past states, since it ignores the fact that new observations typically contain information about the past as well as the present. This is disadvantageous both because knowledge of past states may be intrinsically valuable, and because it impairs learning about fixed or slowly changing parameters of the environment. For these reasons, in offline data analysis it is usual to infer on every set of states using the entire time series of observations, an approach known as (fixed-interval) Bayesian smoothing. Unfortunately, however, this is impractical for real agents, since it requires the maintenance and updating of beliefs about an ever-growing set of states. We propose an intermediate approach, finite retrospective inference (FRI), in which agents perform update beliefs about a limited number of past states. (Formally, this represents online fixed-lag smoothing with a sliding window.) This can be seen as a form of bounded rationality in which agents seek to optimise the accuracy of their beliefs subject to computational and other resource costs. We show through simulation that this approach has the capacity to significantly increase the accuracy of both inference and learning, using a simple variational scheme applied to both randomly generated Hidden Markov models (HMMs), and a specific application of the HMM, in the form of the widely used probabilistic reversal task. Our proposal thus constitutes a theoretical contribution to normative accounts of bounded rationality, which makes testable empirical predictions that can be explored in future work.

## Introduction

To behave adaptively, agents need to continuously update their beliefs about present states of the world using both existing knowledge and incoming sensory information, a process that can be formalised according to the principles of probabilistic inference (Gregory, 1980; von Helmholtz, 1867). This simple insight has generated a large field of inquiry than spans most areas of the mind and brain sciences and seeks to build probabilistic accounts of cognition (Aitchison and Lengyel, 2016; Clark, 2012; Friston, 2010; Pouget et al., 2013; Rao and Ballard, 1999; Tenenbaum et al., 2011).

In this paper, we take this framework for granted, and consider an important and related problem, that of using new sensory information to update beliefs about the past. This is important because, under conditions of uncertainty, new observations can contain significant information about past states as well as present ones (Corlett et al., 2004; FitzGerald et al., 2017; Moran et al., 2019; Shimojo, 2014a).

In offline cognition or data analysis (in which agents are dealing with complete data sets, and are not required to respond to them in real time), it is possible to make inferences about all time points simultaneously (Figure 1).

**Figure 1.**
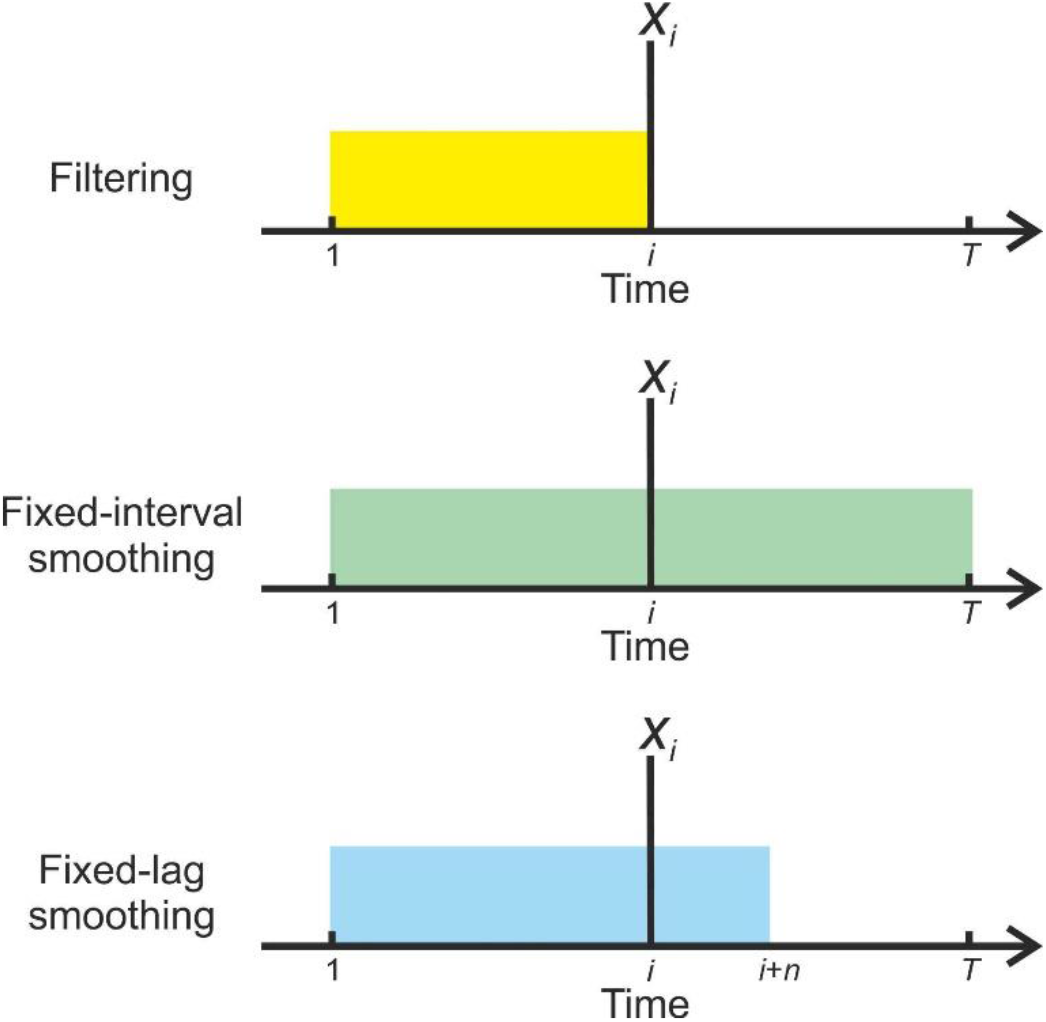
**Illustration of the information used by different strategies for inference when forming beliefs about state** *x* **at time** *i* **(indicated by the vertical line) In filtering (top,) beliefs are based solely on observations made up to and including that time (o**_1:*i*_), **as indicated by the yellow block, and are not revised in the light of subsequent information. In fixed-interval smoothing (middle), observations from the whole data set (o**_1:*T*_**) are used to inform each set of beliefs, as indicated by the green block. In fixed-lag smoothing (bottom), beliefs are retrospectively updated up to some fixed lag** *n*, **so o**_1:*i*+*n*_ **are used (indicated by the blue block). (See Methods for a more formal description, and explanation of the notation) Fixed-lag smoothing allows an agent to perform finite retrospective inference, which constitutes a principled trade-off between the reduced inferential accuracy resulting from filtering and the potentially severe computational costs of retrospection to an indefinite temporal depth.**

In other words, one uses every observation to inform every belief about hidden states. This option is unavailable to real, embodied agents because they need to perceive and act in time (online). They thus need to perform retrospective inference to increase the accuracy of their beliefs about the past. To perform retrospective inference optimally (or, equivalently in this context, to be strictly rational) it is necessary for an agent to update beliefs about a sequence of states stretching backwards to the beginning of the current task or context, or perhaps even to the beginning of its existence. This sequence is both indefinitely long and constantly growing, and representing and updating these beliefs will thus, in many situations, place intolerable demands on any real organism.

We propose an alternative approach, finite retrospective inference (FRI), in which agents update beliefs about states falling within a limited temporal window stretching into the past (FitzGerald et al., 2017). Selecting the size of this window, and thus the depth of retrospective belief updating constitutes a form of bounded rationality (Gigerenzer and Goldstein, 1996; Ortega et al., 2015; Simon, 1972), since it trades off inferential accuracy against resource costs (for example, the metabolic and neuronal costs associated with representing beliefs, and the time to perform the calculations). The depth of updating performed by an agent in a particular context might be selected using a form of ‘metareasoning’ in response to environmental demands (Lieder and Griffiths, 2017; Russell and Wefald, 1991). In particular, it is likely that where observations are noisier, and/or temporal dependencies are greater (in other words, where the past remains significant for longer) such strategies will be more advantageous, and are likely to be favoured, provided that other constraints allow it. Alternatively, the degree of retrospection might be phenotypically specified (and thus, presumably, selected for during species evolution). In either case, a bounded-rational approach to retrospection has the potential to explain and quantify how humans and other organisms approach but do not attain optimal performance on a number of cognitive tasks.

FRI differs from existing probabilistic accounts of online cognition (Aitchison and Lengyel, 2016; Friston and Kiebel, 2009; Glaze et al., 2015; Ma et al., 2006; Rao and Ballard, 1999), which typically only consider inferences about present states, an approach known as ‘Bayesian filtering’ (though see (Baker et al., 2017; Friston et al., 2017; Kaplan and Friston, 2018; Rao et al., 2001)). It thus constitutes a novel hypothesis about cognitive function that extends probabilistic models to subsume a broader range of problems. Importantly, as we will illustrate in the simulations described below, FRI makes testable predictions about behaviour and brain activity in real agents that can be tested in future experimental studies.

## Materials and Methods

### Approximating normative inference

Consider the situation in which an agent seeks to infer on a series of *T* time-varying hidden states **x**_1:*T*_ = (**X**_1_…,**X**_*T*_} given a set of time-invariant parameters **θ** that are known with certainty, a series of observations **o**_1:*T*_ = {**o**_1_,…,**o**_*T*_}, and an initial distribution **x**_0_. To simplify our discussion, in what follows we will assume that all the processes under consideration share the following conditional independence properties:

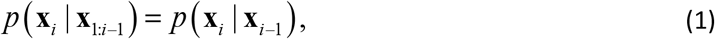

meaning that states at time *i* depend only upon the states at the immediately preceding time (this is the Markov property) and that

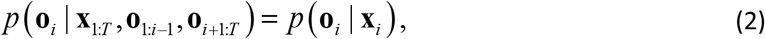

meaning that observations depend only on the current states, and not previous states or observations. However, the general principles presented in this paper apply equally in cases where many, if not all, of these properties are relaxed (for example in processes with a higher-order temporal structure). (We start with a general discussion of retrospective inference, which makes no specification about the nature of states and observations, before discussing a specific instantiation below (the HMM)).

By the chain rule of probability, the joint distribution over all states is given by

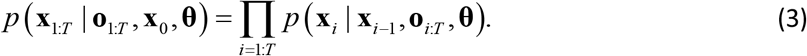

However, inferring on the joint distribution rapidly becomes computationally intractable, and is often unnecessary. Thus instead of inferring on the joint distribution we can instead infer on the marginal distributions over states at each time point (infer on the sequence of most likely states rather than the most likely sequence of states), an approach known as fixed-interval Bayesian smoothing (Sarkka, 2013):

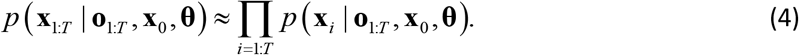

This provides a powerful approach for analysing sequential data, and is widely used in offline data analysis. However, it presents serious practical difficulties for agents performing online inference of the kind that is mandatory for real, embodied agents. (That is, where agents have to make inferences, and very likely take actions, whilst the process is unfolding). Specifically, it requires the agent to store and update an ever-growing set of beliefs about the past, resulting in a set of calculations that will rapidly overwhelm the cognitive capacities of plausible embodied agents. This means that ‘true’ rationality, defined here as cognition that accords precisely with the principles of optimal probabilistic inference, is impossible for real agents, who must instead seek a feasible approximation.

In probabilistic models of online cognition, this is typically achieved by conditioning inference only on past and current observations, an approach known as Bayesian filtering (Sarkka, 2013). This means that agents make inferences of the form

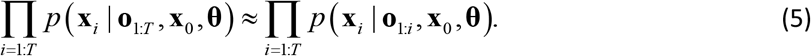

This can be implemented in a straightforward fashion by recursive application of

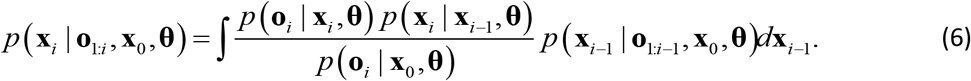

Filtering is thus computationally parsimonious, since it requires only a single set of calculations at each time step and only requires an agent to store fixed beliefs about the past (in the case of first-order processes, only about the immediately preceding time step). However, this parsimony comes at a cost, since it reduces the accuracy of an agent’s beliefs about the past, and consequently, as will discussed later, impairs learning.

To remedy this, an agent that is performing Bayesian filtering, can implement smoothing recursively by performing an additional ‘backwards pass’ through the data:

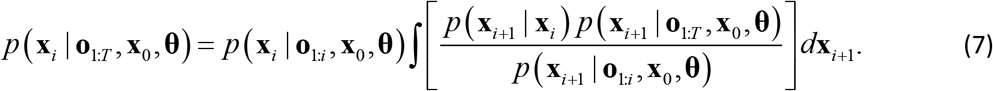

(Use of an integral here and in Equations 8 and 10 presupposes that states are continuous-valued. In the case of discrete states, as in the HMM discussed below, this is replaces with a summation). Here *p*(**x**_*i*_ | **o**_1:*i*_, **x**_0_, **θ**) is the state estimate derived from filtering, *p*(**x**_*i*+1_ | **x**_*i*_) is the dynamic model governing transitions between states, *p*(**x**^*i*+1^| **0**_1:*T*_, **x**_0_, **θ**) is the smoothed state estimate at *i* + 1, and *p* (**x**_*i*+1_| **0**_1:*i*_, **x**_0_, **θ**) is the predicted distribution at *i* +1 given by

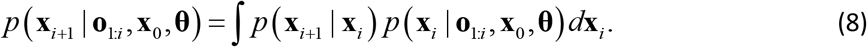

Thus (fixed-interval) smoothing can be carried out in a straightforward manner, beginning with the current state estimate derived from filtering, and working iteratively backwards. Nonetheless, it requires the agent to perform a set of calculations that grows linearly with the time series, and store a similarly growing set of beliefs about past states, and thus introduces significant extra costs for an agent over and above filtering, which are likely to become unsustainable for real agents in ecological contexts. We thus propose that agents make use of an intermediate strategy, finite retrospective inference, in which they perform retrospective belief updating to a limited degree, in a manner that reflects both the desirability of accurate inference and the need to limit resource (and other) costs.

### Finite retrospective inference

To implement FRI, we propose that agents perform fixed-lag smoothing, an approach that is intermediate between full (fixed-interval) smoothing and filtering. In fixed-lag smoothing, agents update beliefs about all states within a fixed-length time window that includes the present time but stretches a set distance into the past (Figure 1) (FitzGerald et al., 2017). This window moves forward in time at the same rate that observations are gathered, meaning that cognition occurs within a sliding window. (In principle the sliding window approach can also be used to infer on the joint distribution of short sequences of states (FitzGerald et al., 2017), but we focus on smoothing in this paper for the sake of simplicity) We are unaware of a precedent for this approach in treatments of cognition, however it has been employed in other contexts (Chen and Tugnait, 2001; Cohn et al., 1994; Moore, 1973; Sarkka, 2013). This means that, for a window of length *n* (one that stretches *n* – 1 timesteps into the past) considered at time *t*, agents approximate the true marginal distribution as follows:

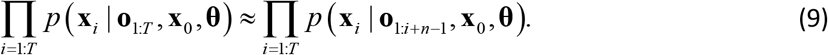

As can be seen by comparing equations (5) and (9), filtering is a special case of fixed-lag smoothing in which *n* = 1. Smoothing can thus be performed by iteratively evaluating

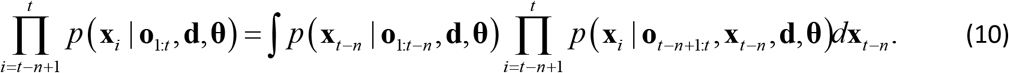

This simply requires the agent to track *p*(**x**_*t–n*_ | **o**_1:*t–n*_, **d, θ**), the filtered estimate of the states that obtain at the timestep immediately preceding the current window. Practically, fixed-lag smoothing can be implemented using equation (7), with the proviso that backward recursion is only performed *n*–1 times. In other words, rather than propagating new information right the way back through a time series as is typical in offline applications, it is only propagated to a fixed depth (*n*–1), limiting the computational cost to the agent. This allows agents to adopt a bounded rational strategy in which they trade off inferential accuracy and computational (and potentially other) costs to select an appropriate depth of processing.

### Parameter learning using retrospective inference

We next consider the more general situation in which there is uncertainty about both states and parameters, and agents must therefore perform learning as well as inference. This is often referred to as a ‘dual estimation’ problem (Friston et al., 2008; Radillo et al., 2017; Wan et al., 1999), and is characteristic of many real-world situations. To do so, we condition beliefs about parameters on a set of hyperparameters **λ**, such that states and observations are independent of the hyperparameters when conditioned on the parameters, meaning that

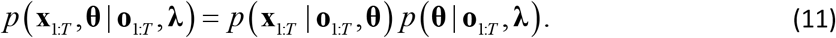

Learning and inference are inextricably related to one another, since beliefs about states depend on beliefs about parameters, and vice versa. Since parameters are fixed, accurately estimating them involves accumulating evidence across entire time series, and thus beliefs about multiple sets of states. This means that increasing the accuracy of beliefs about the past, through retrospective belief updating, also increases the accuracy of parameter estimation. Crucially, since improved parameter estimates will also result in more accurate beliefs about the present and better predictions about the future. Thus, in the context of uncertainty about model parameters, retrospective belief updating is advantageous even for an agent that has no intrinsic interest in the past. This is a very important point, since it argues for the wide importance of retrospective belief updating across a variety of situations and agents.

At a practical level, learning using FRI is very similar to offline learning. We treat each window as a time series in its own right, with **λ** is replaced by 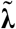, the summary statistics of *p*(**θ** | **o**_1:*t–n*_, **λ**), which is the posterior distribution over the parameters conditioned on all observations preceding the current window, and perform learning and inference as normal. The use of a sliding window does, however introduce a small additional complexity, since successive windows overlap and thus share data points. Thus, if we treated each window as a separate time series we would count each observation multiple times and, as a result, overweight them. To avoid this, when updating 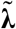 we only use information about states at the first time-point in the window (i.e. *p*(**x**_*t*–*n*+1_| **o**_1:*t*_, **θ**)). (A more specific example of this is provided for the HMM below) This also means that only the best available estimate of each set of states (in other words, the estimate that will not be revised in light of future evidence) contributes to stored beliefs about the parameters of the model.

### Retrospective inference in Hidden Markov models

To illustrate the utility of bounded-rational retrospective inference for an agent, we applied the principles described above to Hidden Markov models (Figure 2). (In principle though, they apply equally to a broad range of models with alternative properties such as continuous state spaces and higher-order temporal structure) In an HMM, the system moves though a series of *T* time-varying hidden states, each of which is drawn from a discrete state space of dimension *K*. Hidden states *x*_1:*T*_ are not observed directly, but instead must be inferred from observed variables. Here we assume that these also discrete, with dimension *M*, but this need not be the case. Thus at time *t* (where 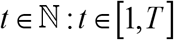), *x_t_* is a binary vector of length *K* such that 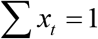 and similarly *o_t_* is a binary vector of length *M* such that 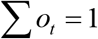. Initial state probabilities are encoded in a row vector **d**, which we will hereafter assume to encode a uniform distribution. Transitions between states are first-order Markovian, and the transition probabilities are encoded in a *K* × *K* matrix **A**, such that

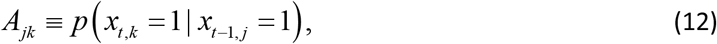

where *A_jk_* ∈ [0,1] and 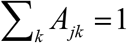. This means that each row **A**_*j*•_, encodes the transition probabilities from state *j* to the entire state space. From this it follows (Bishop, 2006) that

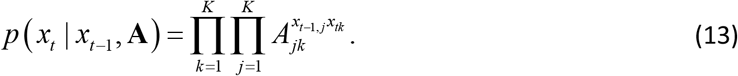

**Figure 2:**
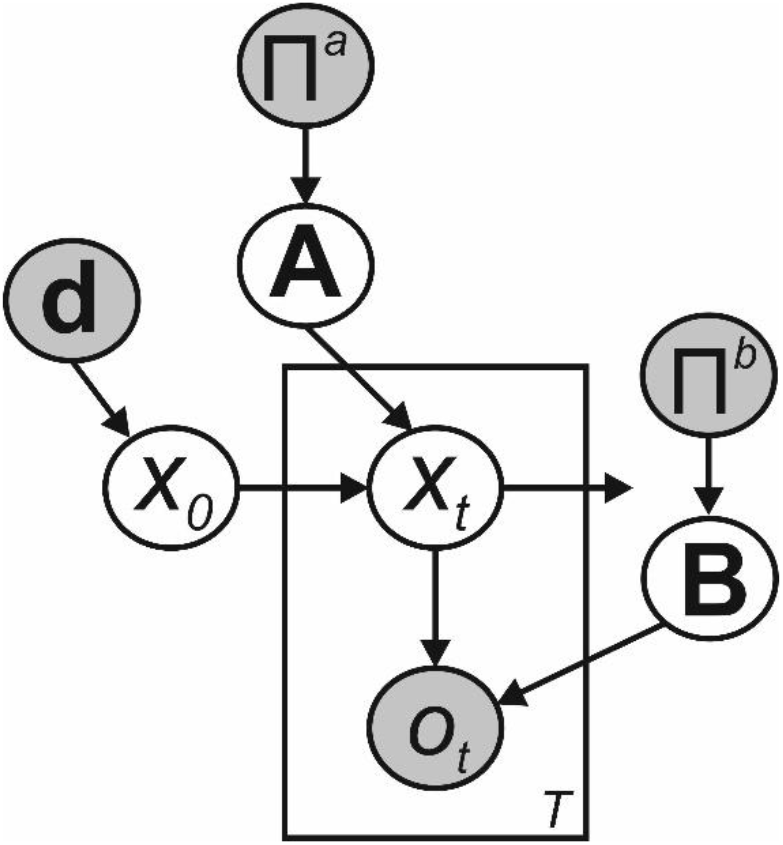
**Bayesian graph illustrating the structure of the Hidden Markov model described in the text. (Shaded circles indicate variables with known values, unshaded circles indicate hidden variables) Transitions between hidden states** *x*_0_ **to** *x_T_* **are governed by the transition matrix A, and are first-order Markovian. Observations** *o*_1_ **to** *o_T_* **depend only on the current hidden state and the emission matrix B. Where the parameters of A and B need to be learnt, as depicted here we include appropriate sets of Dirichlet priors, parameterised by the matrices Π**^a^ **and Π**^*b*^ **respectively. Beliefs about the initial hidden state** *x*_0_ **are governed by the parameter vector d.**

Similarly, the *M*×*K* matrix **B** encodes the emission probabilities such that

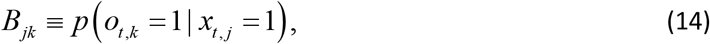

where *B_jk_* ∈ [0,1] and 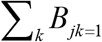. Thus that each row **B**_*j*•_, encodes the probability of each observed variable when in state *j*, and

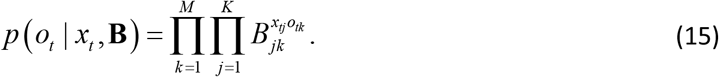

#### Pure inference in HMMs

To calculate the smoothed marginal posterior *γ*(*x_i_*) in an HMM, we can make use of the forward-backward algorithm (Rabiner, 1989). This involves recursive forward and backward sweeps, that calculates two quantities *α*(*x_i_*) and *β*(*x_i_*) for each time point (Bishop, 2006) such that

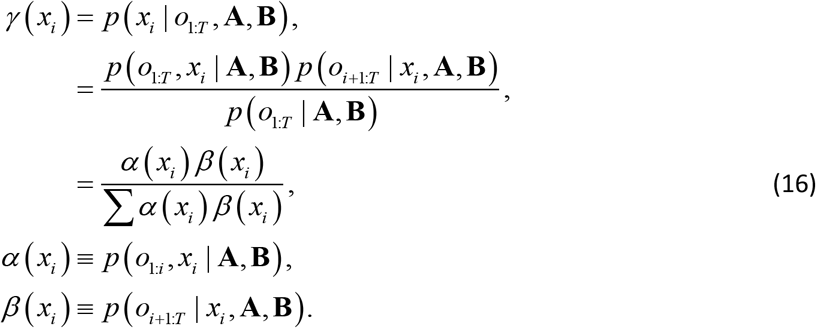

α(*x_i_*) thus corresponds to the unnormalised filtered posterior, and is given by

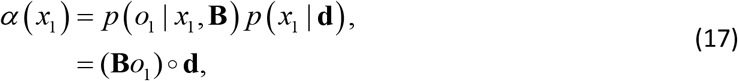

for the first state, and

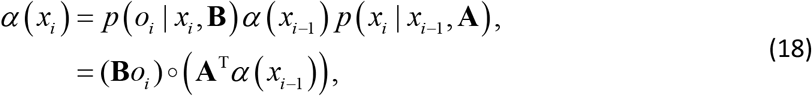

for all subsequent states. Here ∘ denotes the Hadamard or element-wise product. *β*(*x_i_*) is given by

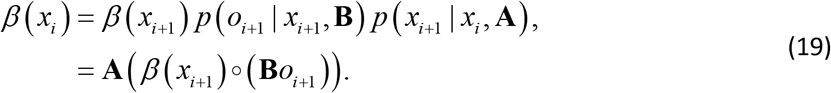

To apply the sliding window approach to this model, at each timestep we simply evaluate the filtered posterior using (18) and then perform backward inference a fixed number of steps using (19).

#### Dual estimation in HMMs

To learn the transition probabilities of an HMM we first need to define an additional quantity, the dual-slice marginal *ξ*(*x_i_, x*_*i*–1_), which corresponds to the joint probability distribution *p*(*x_i_, x*_*i*–1_ | *o*_1:*T*_, *θ*) (Baum et al., 1970; Bishop, 2006). It is simple to show that

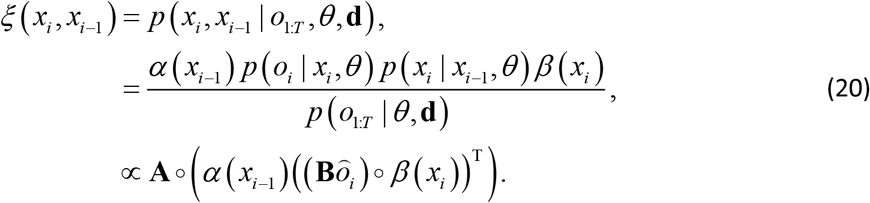

(For a more detailed exposition of this see (Bishop, 2006).)

Introducing learning renders exact inference impossible, which necessitates the use of an approximation. Broadly speaking, such approximations fall into two categories: sampling approaches (Andrieu et al., 2003), which are computationally expensive but asymptotically exact, and variational approaches which are more computationally efficient but require the introduction of a tractable approximate distribution (Blei et al., 2017). We focus here on implementing model inversion using variational Bayes (Beal, 2003), which we believe has some neurobiological plausibility (Friston et al., 2017). This is not a strong claim, however, about the actual mechanisms used by human observers (or indeed any other agent), and similar results could be derived under any appropriate scheme. (See Appendix for further description of the variational methods employed here)

In the offline case, this model has been described in (Beal, 2003; Mackay, 1997), and the reader is referred to these sources for more detailed expositions. Briefly, we start by placing Dirichlet priors over each row of the transition matrix **A** and the observation matrix **B** such that

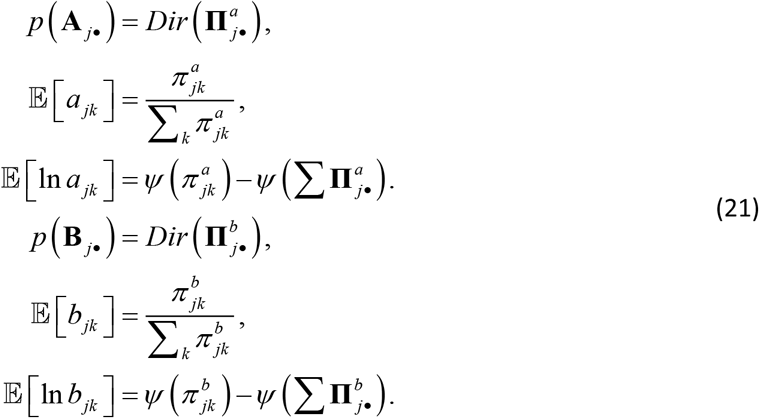

where **Π**^*a*^ and **Π**^*b*^ are matrices encoding the concentration parameters of the Dirichlet distributions, and *ψ* is the digamma function. Since the Dirichlet distribution is the conjugate prior for a multinomial likelihood, this enables us to carry out parameter learning using a set of simple update equations as described below.

The log joint probability distribution for the model thus becomes

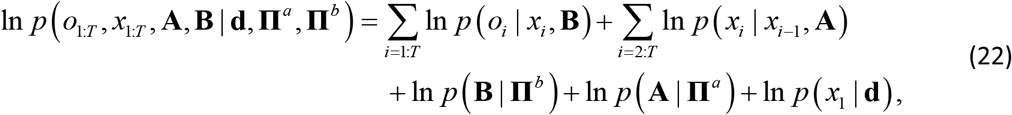

and model inversion can be performed by iteratively evaluating the following update equations for the states and parameters (see Appendix for a full derivation).

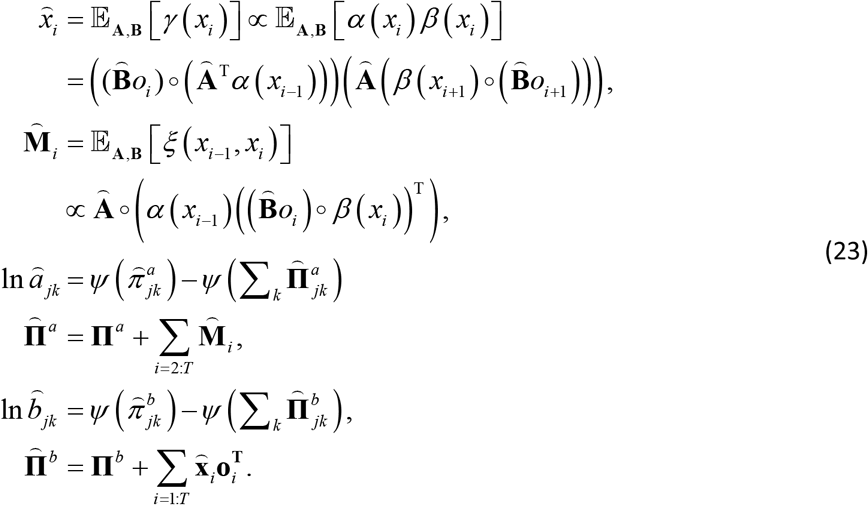

This means that inference about the smoothed *γ*(*x_t_*) and dual-slice marginals *ξ*(*x_t_, x*_*t*−1_) is calculated by applying the forward-backward algorithm at each iteration, using the variational estimates 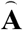 and 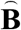, in place of the non-Bayesian **A** and **B** used in Eq 17–19 (Mackay, 1997). The update equations for the parameters also have intuitive interpretations. Updates of the transition matrix **A** correspond to accumulating evidence about the number of times each state transition occurs, whilst those for the observation matrix **B** correspond to a similar evidence accumulation process, this time about the number of times that a particular observation was made whilst occupying a particular state.

For the variational HMM, the lower bound *L* can be calculated in terms of the normalisation constants 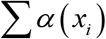 derived during filtering, and the Kullback-Leibler divergences between prior and posterior distributions over the parameters (see (Beal, 2003; Bishop, 2006) for derivationa). Thus

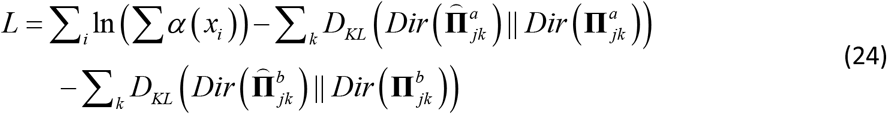

In all simulations, iterations were performed until the difference in the variational lower bound *L* was less than 1^−6^ times the number of data points (*T*). To carry out online learning and inference, we simply apply the sliding window approach described earlier to this model. This means that we only evaluate 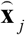 and 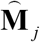 for timepoints that fall within the current window, and parameter learning is performed by updating the concentration parameters using the following equations:

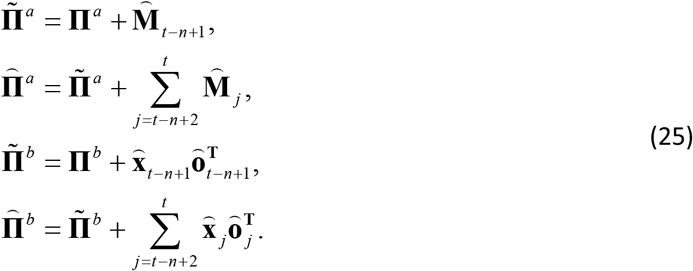

Here 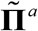 and 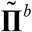 denote the fixed-lag parameters that are incremented across time steps, **Π**^*a*^ and **Π**^*b*^ denote the values of the fixed-lag concentration parameters from the previous time step (in other words, the evidence that has been accumulated prior to the current window), and 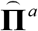 and 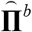 denote the full estimates of the concentration parameters based on timesteps 1 to *t*.

#### The probabilistic reversal task as a special case of the HMM

To illustrate the utility of FRI even for relatively straightforward tasks, we simulated inference and learning on a probabilistic reversal paradigm (Glaze et al., 2015; Hampton et al., 2006; Radillo et al., 2017). Briefly, subjects are required to track an underlying hidden state that occasionally switches between one of two possible values, based on probabilistic feedback (in other words, feedback that is only, for example, 85% reliable) This paradigm is both simple and widely used, and the small state space makes illustrating results in graphical form relatively straightforward. In addition, the fact that the paradigm is widely used makes it an appealing tool for exploring to what extent human subjects actually employ FRI when solving this sort of task. The task can be modelled as an HMM, in which there are only two hidden states *x_t_* ∈ {1,2}, which probabilistically generate one of two possible observations *o_t_* ∈ {1,2} (Costa et al., 2015; FitzGerald et al., 2017; Hampton et al., 2006; Schlagenhauf et al., 2014). The parameter *r* encodes the probability of a reversal between trials, and *v* encodes the reliability of observations. Thus:

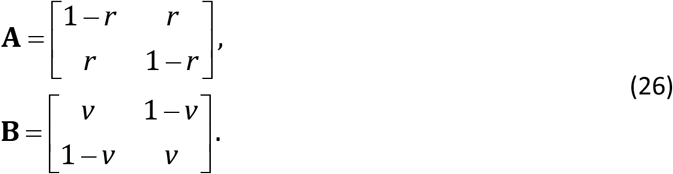

(Introducing learning requires a slight modification of the standard HMM parameter update equations to reflect the symmetry of the **A** and **B** matrices, as described in the Appendix.)

### Simulations

#### Probabilistic reversal task

To illustrate the effects of retrospective inference on an agent’s beliefs whilst doing the probabilistic reversal task, we simulated 1000 instantiations of a 256 trial task session, with parameters set as *r* = 0.1 and *v* = 0.85, plausible values for real versions of the task (e.g. (FitzGerald et al., 2017)). For the ‘pure inference’ agent, we set extremely strong (and accurate) prior beliefs about **A** and **B** by setting initial values of **Π**^(*a*)^ = 10^6^ **A** and **Π**^(*b*)^ = 10^6^**B**. This has the consequence of effectively fixing these parameters to their prior values (in other words, essentially rendering them fixed parameters). For the ‘dual estimation’ agent, we kept the prior beliefs about **B** identical, but set weak priors on the transition matrix of

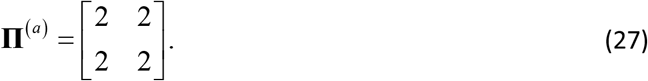

This has the consequence of allowing agents beliefs to be determined almost completely by the data they encounter. Window lengths for retrospective inference were set at 1, 2, 4, 8, 16, 32, 64, 128 and 256 trials, and we also simulated an agent performing offline (fixed interval) smoothing for comparison. To assess the accuracy of inference and learning, we calculated the log likelihood assigned to the true sequence of hidden states and the true value of *r*, calculated using Eq 36 (see Appendix), and averaged these across simulations.

The aim of these simulations is to demonstrate the effects, and potential advantages, of performing FRI for an agent, even on relatively simple tasks. However, establishing whether retrospective inference is in fact a feature of human cognition requires careful experimental validation. This will involve careful model-based analysis of behavioural (and possibly neuroimaging) data collected on appropriate behavioural tasks. We intend to address this in future studies.

#### Random HMMs

To show that the effects that we illustrate are not due to some specific feature of the probabilistic reversal paradigms, we performed similar simulations, this time using HMMs with three possible hidden states, three possible observations, and randomly generated transition probabilities. We generated 10 such HMMs, and simulated 100 instantiations of each, whilst varying the diagonal terms of the emission matrix **B** at intervals of 0.05 between 0.65 and 0.95 (and setting the off-diagonal terms to be equal). (This corresponds to varying the degree of perceptual uncertainty) Prior beliefs for the pure inference and dual estimation agents were set as described for the reversal task, and accuracy was assessed in a similar manner, using Eq 23.

## Results

To explore the properties of fixed-lag retrospection in pure inference problems (in other words, ones where no learning is necessary), we simulated behaviour on both the probabilistic reversal task and on random HMMs. As expected, in both cases, FRI considerably improved the accuracy of agents’ final (offline) beliefs about past hidden states. (Online estimates of current states are identical under all approaches). Strikingly, in both cases, this improvement occurred even when agents only retrospected over short windows (Figure 3), suggesting that, in certain problems at least, a limited capacity for retrospection can yield significantly improves inference.

**Figure 3:**
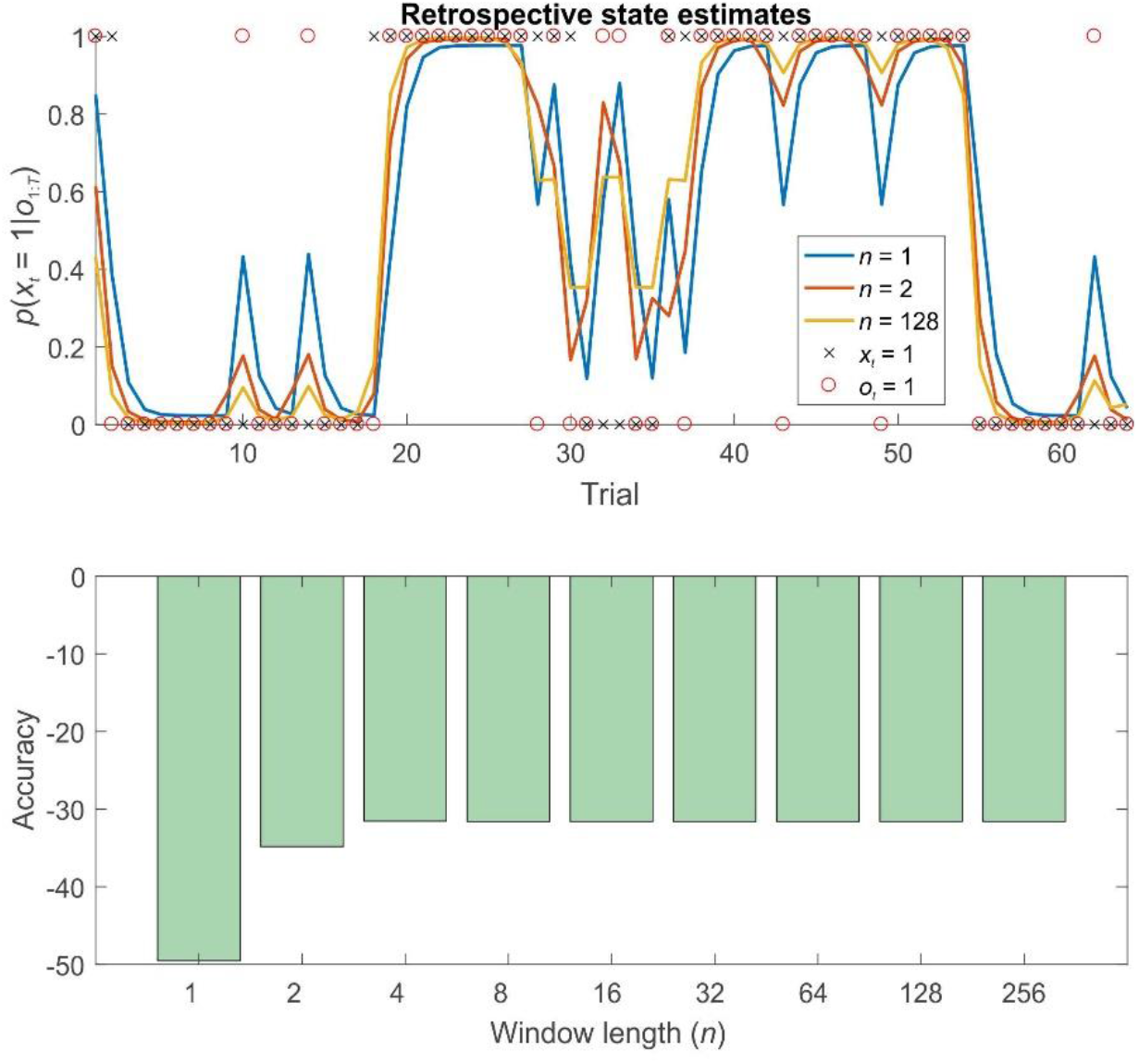
Retrospective belief updating improves state estimates during pure inference on a probabilistic reversal task. Top panel: illustration of the first 64 trials of a 256 trial session of the reversal task using different strategies. The final (retrospective) posteriors are shown in blue (*n*=1, filtering), orange (*n*=2) and gold (*n*=128). Black crosses show the true hidden state, and red circles the observations made on each trial. Retrospective belief updating allows agents to infer the true underlying states more accurately. Bottom panel: relative log accuracy of models of different window lengths, averaged across simulated time series (see main text for details). (Accuracy is quantified as the log likelihood assigned to the true sequence of states by the agent, averaged across simulations) This illustrates that, in this context at least, even the use of a very short window leads to significantly more accurate beliefs, but that this benefit saturates relatively rapidly (by about *n*=8). Thus for a bounded rational agent performing pure inference, the optimal window length may be surprisingly low, depending on the relevant computational costs.

Simulations of dual estimation problems in which there was uncertainty about *r* clearly illustrated that retrospective inference increases the accuracy of both retrospective *and* online state estimation, as a result of increased accuracy in parameter learning (Figures 4 and 5). One important feature to note is that even when the maximum possible depth of retrospection is employed (*n* = 256), the accuracy of online state estimation always falls significantly short of offline estimation. This indicates the fact that, however great the representational and computational sophistication of an agent, there is always a cost to performing inference online, rather than with a complete data set. If sufficiently high, this cost provides an incentive to perform additional (subsequent) offline processing, perhaps during sleep, and it is conceivable that this might be linked to the extended process of memory consolidation. Similar patterns were observed in the random HMM simulations, supporting the notion that these are general properties of retrospective inference (Figure 6).

**Figure 4:**
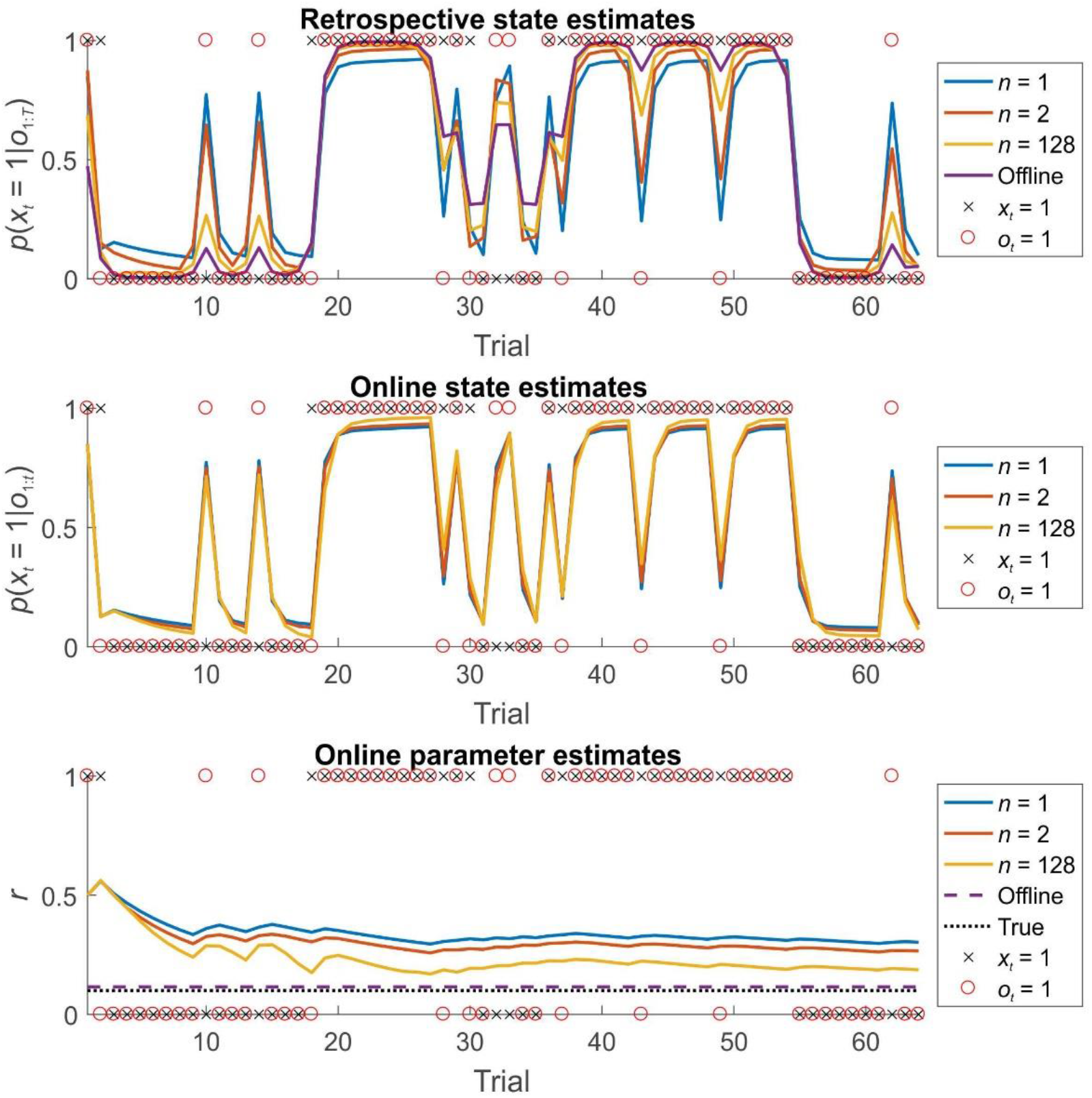
**State and parameter estimation for agents performing dual estimation on the first 64 trials of a 256 trial session of the reversal task using different strategies. Top panel: the accuracy of retrospective belief estimates** *p*(*x_i_* | *o*_1:*T*_) **increases with greater window lengths, but still falls well short of the performance of an offline agents, who has access to the entire time series simultaneously. Middle panel: the accuracy of online (filtered) beliefs about the current state** *p* (*x* | *o*_1:*i*_) **subtly but consistently increases with greater window length. Note that this effect is entirely due to the beneficial effects of greater window lengths on parameter learning. Bottom panel: the effect of window length on parameter learning. Estimates of** *r* **are derived from 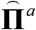 at each timestep (the best estimate available to the agent at that time). With greater window lengths, parameter estimates converge more rapidly on the true value. (Estimates from agents performing retrospective inference with windows of length 1, 2 and 128 time steps are shown in blue, orange and gold respectively. Estimates from an agent performing offline inference is shown in purple. True hidden states are indicated with black crosses, whilst observations are indicated with red circles. The true value of parameter** *r* **is indicated with a dotted black line in 5c)**

**Figure 5:**
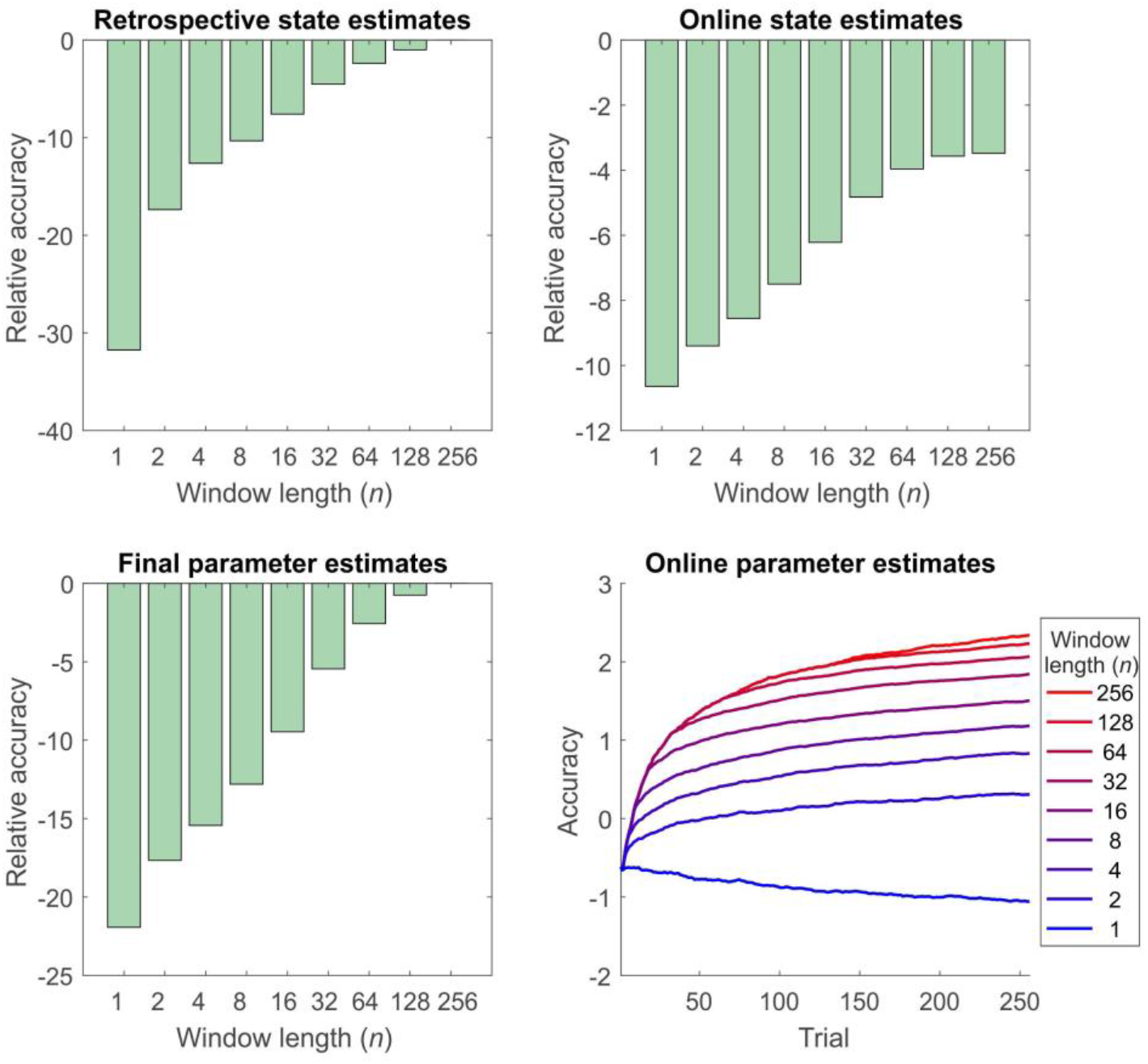
**Accuracy of inference and learning on the reversal task for agents using different window lengths, averaged across 1000 simulations (see ‘Simulations’ for more details). Accuracy is quantified as the log likelihood assigned to the true sequence of states or the true parameter value by the agent, averaged across simulations. Top left panel: accuracy of retrospective state estimation relative to the performance of an offline agent. Accuracy increases with window length, becoming identical for online and offline agents with the same effective window length (256 trials). Top right panel: accuracy of online (filtered) state estimates relative to the filtered state estimates of an offline agent. Accuracy increases with window length, but never becomes equivalent to that of an offline agent. This difference reflects the fact that the parameter estimates of the online agent only use observations made up to the present time, rather than on the entire data set. (In other words,** *p*(*θ*|*o*_1:*t*_, *λ*) **at trial** *t* **rather than** *p* (*θ* | *o*_1:*T*_, *λ*)). **This can be thought of as a cost of online inference. Bottom left panel: accuracy of final parameter estimates relative to the performance of an offline agent. Accuracy progressively increases with window length, becoming equivalent for online and offline agents with the same effective window length. Bottom right panel: average accuracy of parameter estimates across trials. Accuracy of parameter estimation increases with window length, and these differences progressively appear as the session goes on. (Absolute values of the accuracy measure are difficult to interpret here, but the relative accuracy of the difference agents is meaningful).**

**Figure 6:**
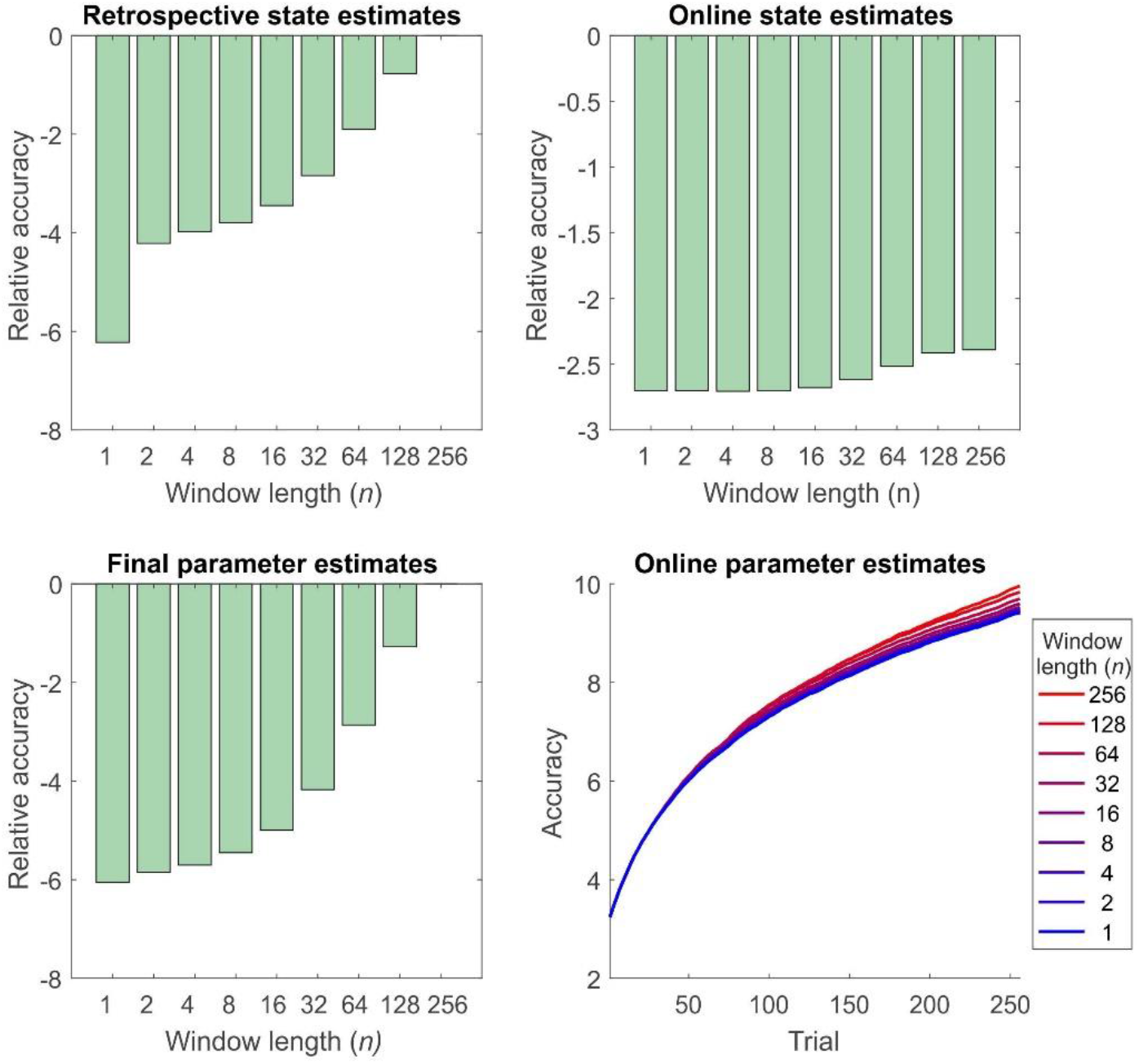
Accuracy of inference and learning for random HMMs (see ‘Simulations’ for more details). In general these mirror the results for the reversal task (Figure 5), but the quantitative differences are smaller, perhaps reflecting the greater number of states and parameters to be estimated. (Accuracy is quantified as the log likelihood assigned to the true sequence of states or the true parameter value by the agent, averaged across simulations) Top left panel: accuracy of state estimation increases with window length, becoming identical for online and offline agents with the same effective window length (256 trials). Top right panel: accuracy of online (filtered) state estimates relative to the filtered state estimates of an offline agent. Accuracy increases with window length, but never becomes equivalent to that of an offline agent. Bottom left panel: accuracy of final parameter estimates relative to the performance of an offline agent. As the window length employed increases, so does accuracy, becoming equivalent for online and offline agents with the same effective window length. Bottom right panel: average accuracy of parameter estimates across trials. Accuracy of parameter estimation increases with window length, and these differences progressively appear as the session goes on.

## Discussion

In this paper, we consider the problem of accurately updating beliefs about the past from the perspective of probabilistic cognition. Specifically, we propose that humans and other agents use finite retrospective inference, in which beliefs about past states are modifiable within a certain temporal window, but are fixed thereafter. We show, using simulations of inference and learning in the context of a probabilistic reversal task, that even a fairly limited degree of retrospection results in significantly improved accuracy of beliefs about both states and parameters. Importantly, the hypothesis that agents perform retrospective inference makes clear predictions about behaviour on appropriate tasks that are quantitatively dissociable from those made under the hypothesis that agents use pure filtering. Implementing retrospective inference also makes specific predictions about brain function, since it requires beliefs about past states to be explicitly represented and updated. Our work thus provides testable hypotheses that can be explored in future behavioural and neurobiological studies.

Perhaps the most significant feature of our simulations is the demonstration that, where there is uncertainty about time-invariant model parameters, finite retrospective inference significantly improves the accuracy of learning. This is important both because these parameters may be of intrinsic interest, and because better learning will result in more accurate beliefs about present and future states. Even if an agent has no intrinsic interest in past events, it still has a clear incentive to perform retrospective inference, since this will allow it to act better in the future. This provides a new twist on the often-advanced hypothesis that the primary function of memory in general, and episodic memory in particular, is to improve predictions about the future (Schacter et al., 2012). Here, in addition to playing a role in constructing imagined future scenarios (Hassabis et al., 2007), the explicit representation of events or episodes in the past may be essential for updating beliefs about current and future states or learning time-invariant properties of an agent’s environment (Baker et al., 2017).

A similar point may be made about the potential importance of retrospective inference for the generation and selection of appropriate cognitive models, a process known as structure learning (Acuña and Schrater, 2010; Braun et al., 2010; Tervo et al., 2016). In this paper, we confine ourselves to considering inference about hidden states and learning about fixed model parameters, but structure learning is an equally important process, and one that is likely to be strongly affected by the depth of retrospective processing employed by an individual. In future work, we plan to address this explicitly, both through simulations and experimental work.

The specific retrospective inference model we describe here differs importantly from previous approaches to modelling probabilistic reversal tasks (Hampton et al., 2006) and change point detection more generally (Radillo et al., 2017; Wilson et al., 2010) in two key ways, first through the fact that we allow for parameter learning (though see (Radillo et al., 2017)), and second, because we simulate agents that are able to update beliefs about past states. Both these processes are important for normative behaviour, and it will be important to establish how closely human performance across a number of domains reflects this. Retrospective inference has also been considered in the context of reinforcement learning (Moran et al., 2019), and we will explore how to approach similar reward learning problems using our probabilistic framework in future. Similar ideas have also been explored in the context of active inference and planning (Friston et al., 2017; Kaplan and Friston, 2018), although these have not explored effects on learning.

Our approach differs importantly from models such as the hierarchical Gaussian filter (Mathys et al., 2011), which use higher-level variables operating at longer time scales to provide an implicit time window, but do not make postdictive inferences of the sort discussed here. (In fact, retrospective inference has the potential to improve accuracy on tasks involving tracking of higher order variables like volatility (Behrens et al., 2007; Mathys et al., 2011), which is a promising area for future study). A closer analogy can be drawn with generalised filtering approaches (Friston, 2008), which infer both on the current state and its derivatives (rate of change, acceleration and so on), and require a finite window of data to perform updates. This similarity is something we intend to return to in future work. Retrospective inference provides a natural explanation for a number of ‘postdictive’ phenomena in perception, in which perception of an event is influenced by other events that only occur afterwards (Eagleman and Sejnowski, 2000; Shimojo, 2014b). A classic example of this is the colour phi phenomenon (Kolers and von Grünau, 1976). Here, two differently coloured dots are briefly displayed to the subject at different spatial locations. If the interval between the flashes is sufficiently short, subjects report perceiving a single moving dot, rather than two separate dots. Critically, they also perceive the colour of the dot as changing during motion, meaning that they perceive the second colour as occurring before it is presented on screen. This means that information about the colour of the second dot has somehow been propagated backwards in (perceptual) time. That such postdictive phenomena might be explained by smoothing has previously been pointed out by (Rao et al., 2001), but our proposal builds on this by suggesting a limited window of updating, as well as highlighting the importance of such belief updating for learning.

The existence of postdictive perceptual phenomena (among other considerations) have led to what is often called the ‘multiple drafts’ account of consciousness (Dennett and Kinsbourne, 1992), in which the contents of conscious are subject to continual revision in the light of new information (at short timescales, at least), and what subjects report is critically dependent upon when they are asked. For example, in the colour phi experiment, subjects’ reported perceptual experience would differ if they were asked to report it before the second dot is shown, as opposed to when they are asked to report it afterwards. This accords extremely well with our proposal (at least if we make the further supposition that the contents of consciousness can, in some sense, be identified with the outcome of optimal perceptual inference). Under FRI (unlike filtering), reported perceptual experience will be critically dependent on when the report is made, since online retrospective inference makes beliefs time-dependent. In other words, my belief about what happened at time *t* may be different depending on whether you ask me for it at time *t* + 1 or time *t* + 10. This means that FRI has the potential to provide the computational underpinning of a ‘probabilistic multiple drafts’ model of perceptual experience.

One intriguing possibility raised by FRI is that different individuals might perform retrospective belief updating to different extents, either on particular tasks or in general, and that this might partially explain between-subject differences in performance on particular tasks (see (FitzGerald et al., 2017) for evidence of this). Such differences might even help explain facets of psychopathology (Montague et al., 2012). For example, impaired learning due to reduced or absent retrospection might lead to the tendency to form delusional beliefs (Adams et al., 2013; Corlett et al., 2004; Hemsley and Garety, 1986). For example, say someone looked at you in an unusual way – making you feel they were spying on you – but then subsequently ignored you: if you could not use the latter information to revise your initial suspicion, you would be more likely to become paranoid about that person. This idea is supported by the finding of altered neuronal responses in subjects with delusions (as compared with healthy controls) during performance of a retrospective belief updating task (Corlett et al., 2007), and is something we intend to return to in future.

Implementing retrospective inference also has important implications for neurobiology. In particular, since agents need to be able to dynamically update beliefs about past states, they are required to store explicit, ordered representations of the past, and it should be possible to find evidence of this in appropriate neuronal structures (Pezzulo et al., 2014). (For some evidence of this, see (Corlett et al., 2004).) Intriguingly, this fits extremely well with an extensive literature on hippocampal function (Fortin et al., 2002; Jensen and Lisman, 2005; Lehn et al., 2009; Pastalkova et al., 2008; Penny et al., 2013), a finding supported by the results of our previous study, which found a relation between depth of retrospective processing and grey matter density in the hippocampus (FitzGerald et al., 2017). On the further supposition that retrospective inference is implemented using filtering and smoothing as described above, this leads to the hypothesis that forward and backward sweeps through recently encountered states, as are known to occur in the hippocampus (Davidson et al., 2009; Diba and Buzsáki, 2007; Pastalkova et al., 2008; Wikenheiser and Redish, 2013) may play a key role in retrospective belief updating. What is less clear, at present, is how to implement retrospective inference within established, neurobiologically-grounded accounts of probabilistic inference in the brain (Aitchison and Lengyel, 2016; Friston, 2005; Ma et al., 2006) – though see (Friston et al., 2017) for related suggestions. This is an extremely important question, and one we intend to return to in future work.

Probabilistic models of cognition are an enormously exciting tool for understanding the complex workings of the mind and brain (Aitchison and Lengyel, 2016; Clark, 2012; Friston et al., 2013; Pouget et al., 2013). The ideas we propose represent a development of such approaches to encompass inference about states in the past, as well as the present. On the further hypothesis that the depth of processing employed is flexible and tailored to the demands of a particular problem or environment, such retrospective processing can also be linked to broader notions of bounded rationality (Gigerenzer and Goldstein, 1996; Ortega et al., 2015; Simon, 1972). We show, through simulations of simple environments, that even a limited degree of retrospection can yield significantly more accurate beliefs about both time-varying states and time-invariant parameters, and thus has the potential to support more adaptive, successful behaviour to justify its extra resource costs. This makes it a plausible strategy for real, biological agents to employ FRI makes both behavioural and neuronal predictions in a number of contexts and thus naturally suggests further avenues for exploration in future work.

## Acknowledgements

THBF is supported by a European Research Council (ERC) Starting Grant under the Horizon 2020 program (Grant Agreement 804701). This manuscript has been released as a Pre-Print at https://www.biorxiv.org/content/10.1101/569574v2.

## Author contributions statement

THBF conceived the study, created the models and performed the simulations. WDP contributed ideas and code. HMB and RA contributed ideas. All authors contributed to writing the manuscript.

## Appendix Derivation of the update equations for the Bayesian HMM

The log joint for the Bayesian HMM we describe is:

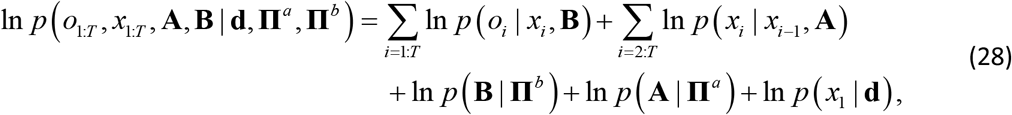

with individual factors given by:

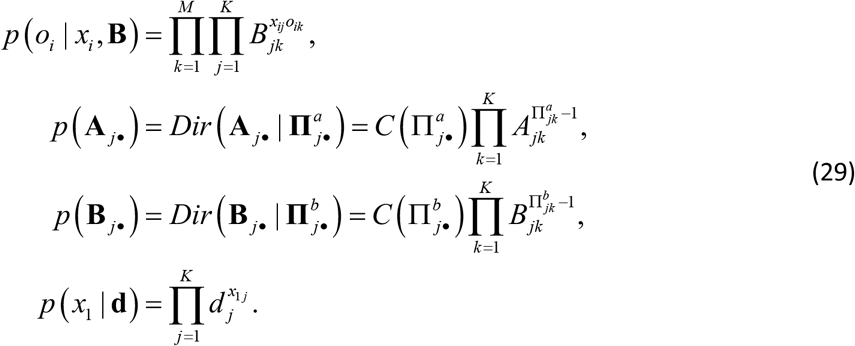

where

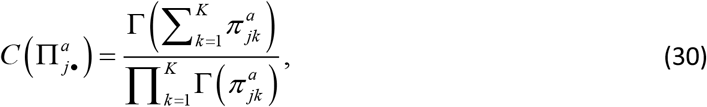

and Γ denotes the gamma function.

Since exact inference in this model is intractable, we define a tractable approximating distribution by factorising such that

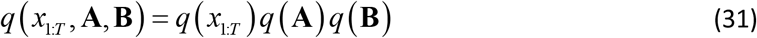

The optimal solution for *q*^*^(*x*_1:*T*_) is given by:

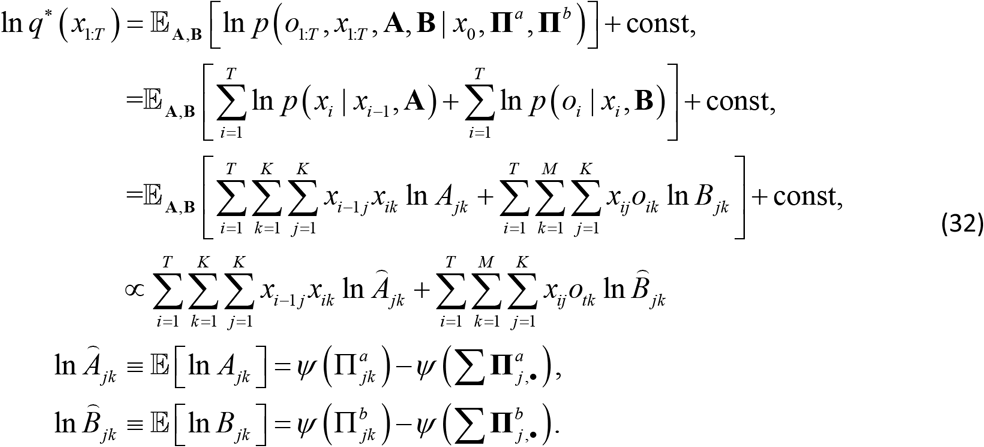

This involves evaluating the entire joint probability distribution over *x*_1:*T*_, which as discussed in the main text rapidly becomes computationally infeasible, and is often unnecessary. However, to infer on the optimal marginal distribution over states, we simply need to perform smoothing using the matrices 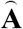 and 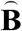. (These are not true transition and observation matrices since the rows do not sum to one. However, the relevant normalisation constants are estimated and applied automatically during smoothing) We can then use the results of smoothing to calculate the joint distributions *ξ*(*x*_*i*–1_, *x_i_*). If we now define a set of summary statistics:

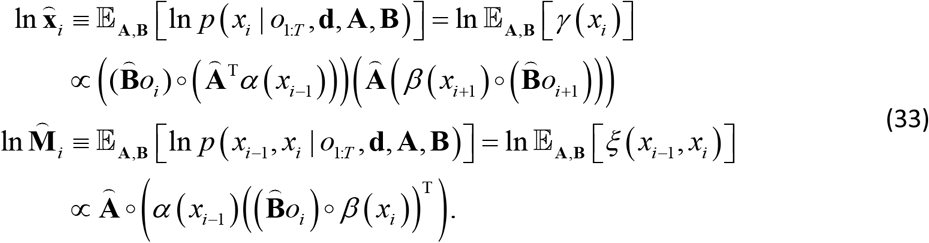

it is easy to find the optimal solutions for *q*^*^(**A**) and *q*^*^(**B**).

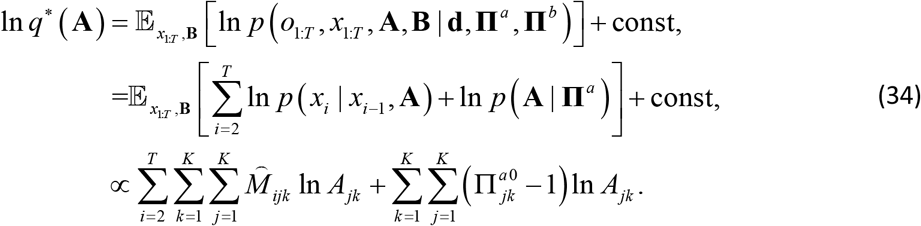

Thus by inspection the optimised factor is:

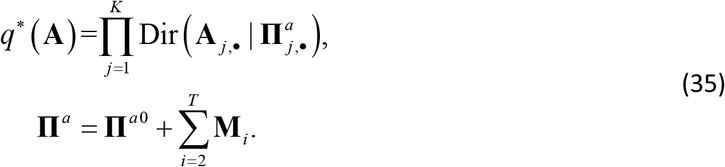

Similarly

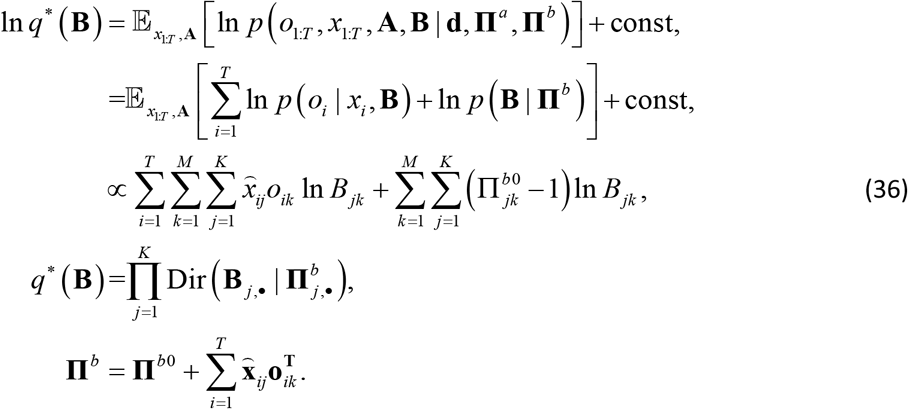

Thus we can perform approximate inference simply by iteratively evaluating the update equations for each factor.

For the probabilistic reversal model, we enforce symmetry between the rows of **A**, and make use of a single Dirichlet prior that has two concentration parameters (in other words, a beta distribution).

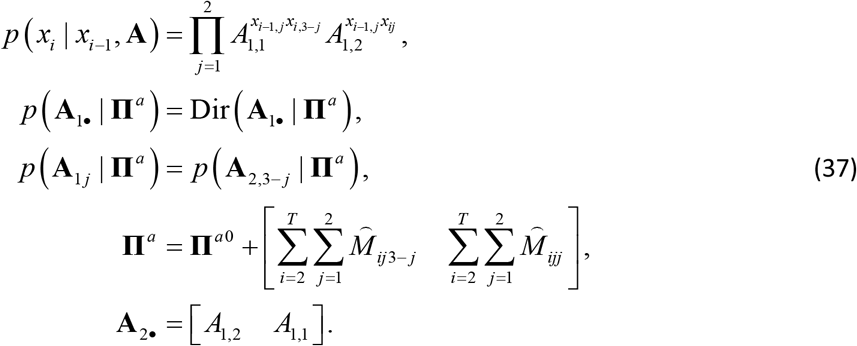

